# Transcriptome-wide characterization of piRNAs during the developmental process of European honey bee larval guts

**DOI:** 10.1101/2022.09.08.507214

**Authors:** Ya-Jing Xu, Qi Long, Xiao-Xue Fan, Ya-Ping Ye, Kai-Yao Zhang, Jia-Xin Zhang, Hao-Dong Zhao, Yu-Tong Yao, Ting Ji, Zhe-Guang Lin, Zhong-Min Fu, Da-Fu Chen, Rui Guo

## Abstract

piRNAs play pivotal roles in suppressing transposons, maintaining genome stability, regulating gene expression, and modulating development and immunity. However, there are few piRNA-associated studies on honey bee, and the regulatory role of piRNAs in the development of bee guts is largely unknown. In this current work, the differential expression pattern of piRNAs during the developmental process of the European honey bee (*Apis mellifera*) was analyzed, and target prediction of differentially expressed piRNAs (DEpiRNAs) was then conducted, followed by investigation of regulatory networks and the potential function of DEpiRNAs in regulating larval gut development. Here, a total of 843 piRNAs were identified in the larval guts of *A. mellifera*; among these, 764 piRNAs were shared by 4- (Am4 group), 5- (Am5 group), and 6-day-old (Am6 group) larval guts, while 11, 67, and 1, respectively, were unique. The first base of piRNAs in each group had a cytosine (C) bias. Additionally, 61 up-regulated and 17 down-regulated piRNAs were identified in the Am4 vs. Am5 comparison group, further targeting 9, 983 genes, which were involved in 50 GO terms and 142 pathways, while two up-regulated and five down-regulated piRNAs were detected in the Am5 vs. Am6 comparison group, further targeting 1, 936 genes, which were engaged in 41 functional terms and 101 pathways. piR-ame-742536 and piR-ame-856650 in the Am4 vs. Am5 comparison group as well as piR-ame-592661 and piR-ame-31653 in the Am5 vs. Am6 comparison group were found to link to the highest number of targets. Further analysis indicated that targets of DEpiRNAs in these two comparison groups were seven development-associated signaling pathways such as Hippo and Notch signaling pathways, seven immune-associated pathways including endocytosis and the Jak/STAT signaling pathway, and three energy metabolism pathways, namely sulfur metabolism, nitrogen metabolism, and oxidative phosphorylation. Moreover, the expression trends of five randomly selected DEpiRNAs were verified based on stem-loop RT-PCR and RT-qPCR. These results were suggestive of the overall alteration of piRNAs during the larval developmental process and demonstrated that DEpiRNAs potentially modulate development-, immune-, and energy metabolism-associated pathways by regulating expression of corresponding genes via target binding, further affecting the development of *A. mellifera* larval guts. Our data offer novel insights into the development of bee guts and lay a basis for clarifying the developmental mechanism underlying the larval guts of European honey bee.

## 1. Introduction

Piwi-interacting RNAs (piRNAs) are a kind of small non-coding RNAs (ncRNAs), with a length distribution from 24 to 32 nt [1]. Accumulating evidence has showed that piRNAs play critical roles in suppressing transposons and maintaining genome stability [2,3]. Dissimilar to miRNA, piRNAs are transcribed from single-stranded RNA (ssRNA) via a dicer-independent mechanism and function by interacting with P-element-induced wimpy testis (Piwi) proteins [4,5]. Structurally, piRNAs have no secondary hairpin structures and biased for uridine (U) at the 5′ end and adenosine (A) signature at position 10 [6]. However, piRNA has been proved to specifically bind to target mRNA in a miRNA-like way [7]. There are two major pathways to generate piRNAs: (1) the primary processing pathway and (2) the ping-pong cycle that amplifies secondary piRNAs [8]. In the cytoplasm, piRNA precursors are transcribed from piRNA clusters, which are processed by the endonuclease Zucchini (Zuc) to produce piRNA intermediates, and subsequently piRNAs can be loaded into Piwi or Aubergine (Aub) to form piRNA induce silencing complexes (piRISCs), which are further trimmed by exonuclease and 2’-O-methylation of piRNAs with Hen1 (nascent helix-loop-helix 1) to produce mature piRNAs. Whereas secondary piRNAs are produced only in germ cells by secondary expansion of Argonaute 3 (AGO3) protein with Aub-piRISCs, this process is also known as the ping-pong pathway [6,9,10,11]. Recent studies demonstrated that piRNAs, as newly emerging regulators of gene expression, could have a function in many biological processes such as immune response and development [12,13]. For example, Wang et al. [14] discovered that piRNAs in *Aedes albopictus* were involved in the host antiviral immune response to Dengue virus 2 (DENV-2) infection.

Honey bee are capable of pollination of a large number of wildflowers and crops, thus playing a pivotal part in maintenance of ecological balance and survival of mankind [15]. As one of the most widely distributed subspecies of *Apis mellifera*, *Apis mellifera ligustica* are commercially reared in China and other countries for their great economic value [16]. Insect gut is an essential organ for food digestion, nutrient absorption, and immune defense [17]. Previous works were mainly focused on adult bee gut and intestinal microorganisms [18,19]. However, little progress on the molecular mechanisms regulating development of bee gut has been made to date.

In the previous study, our group conducted deep sequencing of 4-, 5-, and 6-day-old larval guts of *A. m. ligustica* followed by identification and investigation of miRNAs using bioinformatics [20]. Currently, whether and how piRNAs regulate the development of *A. m. ligustica* larval guts are still largely unknowon. Here, to decipher the expression profile of piRNAs during the developmental process of *A. m. ligustica* larval guts, piRNAs in larval guts were identified and validated based on the obtained high-quality sRNA-seq datasets, and differentially expressed piRNAs (DEpiRNAs) were then analyzed followed by target prediction and regulatory network investigation. DEpiRNAs and corresponding target genes associated with development and immune of larval gut are discussed. Findings in the present study will provide a novel insight into the development of honey bee larval gut and a basis for illustration of the piRNA-regulated mechanism underlying gut development.

## 2. Materials and Methods

### 2.1. Bee larvae

*A. m. ligustica* larvae used in this work were obtained from colonies reared in the apiary at the College of Animal Science (College of Bee Science), Fujian Agriculture and Forestry University, Fuzhou City, China.

### 2.2. Source of sRNA-seq data

Four-, five-, and six-day-old larval guts of *A. m. ligustica* (Am4, Am5, and Am6) were previously prepared, followed by cDNA library construction and deep sequencing with small RNA-seq (sRNA-seq) technology, 38, 011, 613; 43, 967, 518 and 39, 523, 034 raw reads were produced from Am4, Am5, and Am6 groups, respectively, and after quality control, 32, 524, 933; 36, 113, 035 and 27, 691, 488 clean tags were gained. The Pearson correlation coefficients between different biological replicas within each group were above 98.22% [20]. Raw data generated from sRNA-seq were deposited in the NCBI SRA database under the BioProject number.

### 2.3. Identification and investigation of piRNAs

*A. m. ligustia* piRNAs were identified according to our previously described protocol: (1) The clean reads were mapped to the reference genome of *A. mellifera* (Assembly Amel_4.5), and the mapped clean reads were further aligned against GeneBank and Rfam (11.0) databases to remove small ncRNA including rRNA, scRNA, snoRNA, snRNA and tRNA; (2) miRNAs were filtered out from the remaining clean reads; (3) sRNAs with a length distribution from 24 to 33 nt were screened out based on the length characteristics of piRNAs, and only those aligned to a unique position were retained as candidate piRNAs. Next, first base bias of piRNAs in each group was summarized on basis of the piRNA prediction result.

### 2.4. Target prediction and analysis of DEpiRNAs

The expression level of each piRNA was normalized to tags per million (TPM) following the formula TPM = T × 10^6^/N (T denotes clean reads of piRNA, N denotes clean reads of total sRNA). On basis of the standard of *P* value ≤ 0.05 and |log_2_(Fold change)| ≥ 1, DEpiRNAs in Am4 vs. Am5 and Am5 vs. Am6 comparison groups were screened out. TargetFinder software was used to predict target genes of DEpiRNAs [21]. The targets were aligned to the GO (https://www.geneontology.org) and KEGG (https://www.genome.jp/kegg/) databases using the BLAST tool to obtain corresponding annotation.

### 2.5. Construction and analysis of regulatory network of DEpiRNAs

Based on the KEGG pathway annotation information, the target genes annotated in development-, immune-, and energy metabolism-associated signaling pathways were further surveyed to construct the regulatory network, which was then visualized with Cytoscape software [22]. The threshold for screening the targeted binding relationship was set as a binding free energy of less than −15 kcal/mol.

### 2.6. Validation of DEpiRNAs by Stem-loop RT-PCR and RT-qPCR

Total RNA from 4-, 5-, and 6-day-old *A. m. ligustica* larval guts were extracted using a FastPure^®^ Cell/Tissue Total RNA Isolation Kit V2 (Vazyme, Nanjing, China). The concentration and purity of RNA were checked with Nanodrop 2000 spectrophotometer (Thermo Fisher, Waltham, MA, USA). Five DEpiRNAs were randomly selected for stem-loop RT-PCR validation, including four (piR-ame-1146560、piR-ame-1183555、piR-ame-387266 and piR-ame-856650) from Am4 vs. Am5 comparison group and one (piR-ame-592661) from Am5 vs. Am6 comparison group. Specific stem-loop primers and forward primers (F) as well as universal reverse primers (R) were designed using DNAMAN software and then synthesized by Sangon Biotech Co., Ltd. (Shanghai, China). According to the instructions of HiScript ^®^ 1st Strand cDNA Synthesis Kit, cDNA was synthesized by reverse transcription using stem-loop primers and used as templates for PCR of DEpiRNA. Reverse transcription was performed using a mixture of random primers and oligo (dT) primers, and the resulting cDNA was used as templates for PCR of the reference gene snRNA U6. The PCR system (20 μL) contained 1 μL of diluted cDNA, 10 μL of PCR mix (Vazyme, Nanjing, China), 1 μL of forward primers, 1 μL of reverse primers, and 7 μL of diethyl pyrocarbonate (DEPC) water. The PCR was conducted on a T100 thermocycler (Bio-Rad, Hercules, CA, USA) under the following conditions: pre-denaturation step at 95 °C for 5 min; 40 amplification cycles of denaturation at 95 °C for 10s, annealing at 55 °C for 30 s, and elongation at 72 °C for 1 min, followed by a final elongation step at 72 °C for 10 min. The amplification products were detected on 1.8% agarose gel electrophoresis with Genecolor (Gene-Bio, Shenzhen, China) staining.

The RT-qPCR was carried out following the protocol of SYBR Green Dye kit (Vazyme, Nanjing, China). The reaction system (20 μL) included 1.3 μL of cDNA, 1 μL of forward primers, 1 μL of reverse primers, 6.7 μL of DEPC water, and 10 μL of SYBR Green Dye. RT-qPCR was conducted on an Applied Biosystems QuantStudio 3 system (Thermo Fisher, Waltham, MA, USA) following the conditions: pre-denaturation step at 95 °C for 5 min, 40 amplification cycles of denaturation at 95 °C for 10 s, annealing at 60 °C for 30 s, and elongation at 72 °C for 15 s, followed by a final elongation step at 72 °C for 10 min. The reaction was performed using an Applied Biosystems QuantStudio 3 Real-Time PCR System (Themo Fisher). All reactions were performed in triplicate. The relative expression of piRNA was calculated using the 2^−ΔΔCt^ method [23]. Detailed information about primers used in this work is shown in **Table S1**.

### 2.7. Statistical analysis

Statistical analyses were conducted with SPSS software (IBM, Amunque, NY, USA) and GraphPad Prism 7.0 software (GraphPad, USA). Data were presented as mean ± standard deviation (SD). Statistics analysis was performed using Student’s *t* test. Significant (*P* < 0.05) GO terms and KEGG pathways were filtered by performing Fisher’s exact test with R software 3.3.1 [24,25].

## 3. Results

### 3.1. Identification and characterization of piRNAs in A. m. ligustica larval guts

A total of 843 piRNAs were identified in the larval guts of *A. m. ligustica*; among these, 764 piRNAs were shared by Am4, Am5, and Am6 groups, while 11, 67, and 1, respectively, were unique. Further investigation showed that the first base of piRNAs in Am4, Am5, and Am6 groups had a C bias (**Figure 1**).

**Figure 1.**
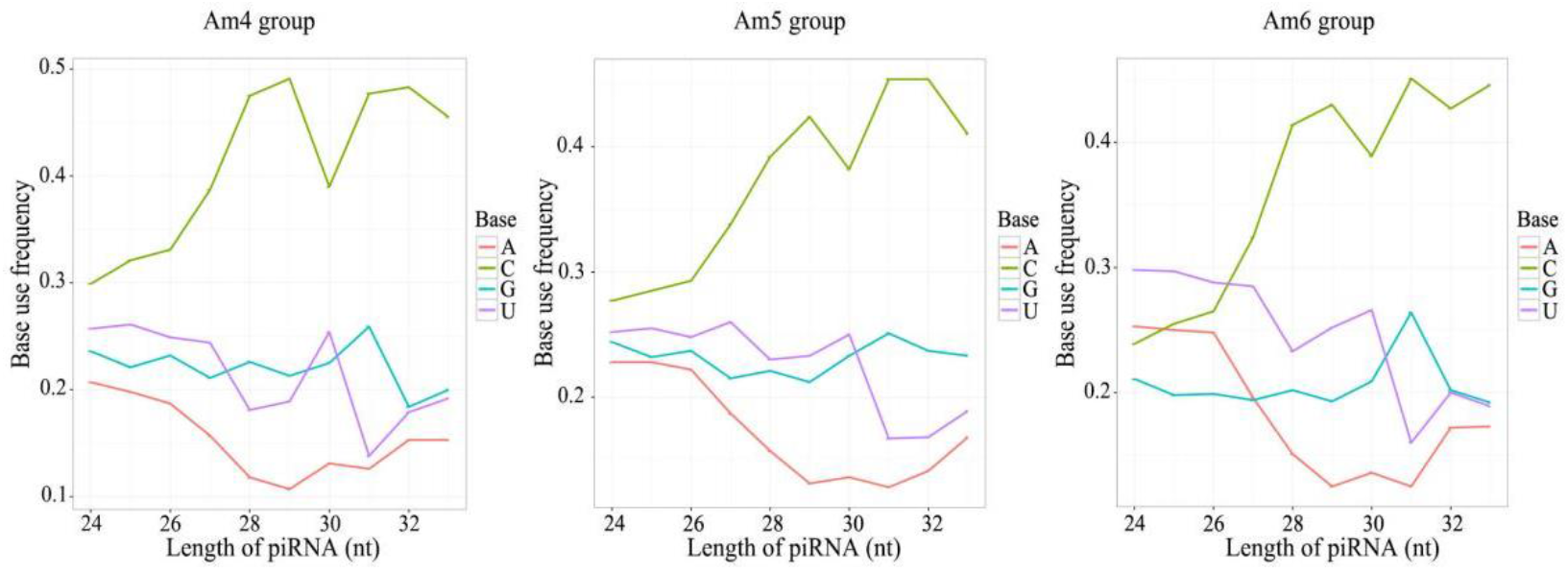
First base bias of piRNAs in Am4, Am5, and Am6 groups.

### 3.2. Differential expression profile of piRNA during the developmental process of larval guts

Here, 78 DEpiRNAs were identified in the Am4 vs. Am5 comparison group, including 61 up-regulated and 17 down-regulated piRNAs; among these, the most significantly up-regulated one was piR-ame-1009988 (log_2_FC=14.52, *P*=7.22E-05), followed by piR-ame-14055 (log_2_FC=14.52, *P*=7.22E-05) and piR-ame-456655 (log_2_FC=14.52, *P*=7.22E-05), while the three most significantly down-regulated DEpiRNAs were piR-ame-1223398 (log_2_FC= −11.38, *P*=3.19E-09), piR-ame-1186994 (log_2_FC= −1.66, *P*=0.041), and piR-ame-1077365 (log_2_FC= −1.65, *P*=0.001) (**Figure 2**A). Seven DEpiRNAs were identified in the Am5 vs. Am6 comparison group, including two up-regulated and five down-regulated piRNAs; among these, the two most significantly up-regulated DEpiRNAs were piR-ame-1243913 (log_2_FC=3.01, *P*=0.019) and piR-ame-592661 (log_2_FC=1.14, *P*=0.005); whereas the most significantly down-regulated piRNAs was piR-ame-1246710 (log_2_FC=−1.18, *P*=0.045), followed by piR-ame-1173337 (log_2_FC=−10.96, *P*=1.34E-05) and piR-ame-31653 (log_2_FC=−10.96, *P*=1.34E-05) (**Figure 2**B).

**Figure 2.**
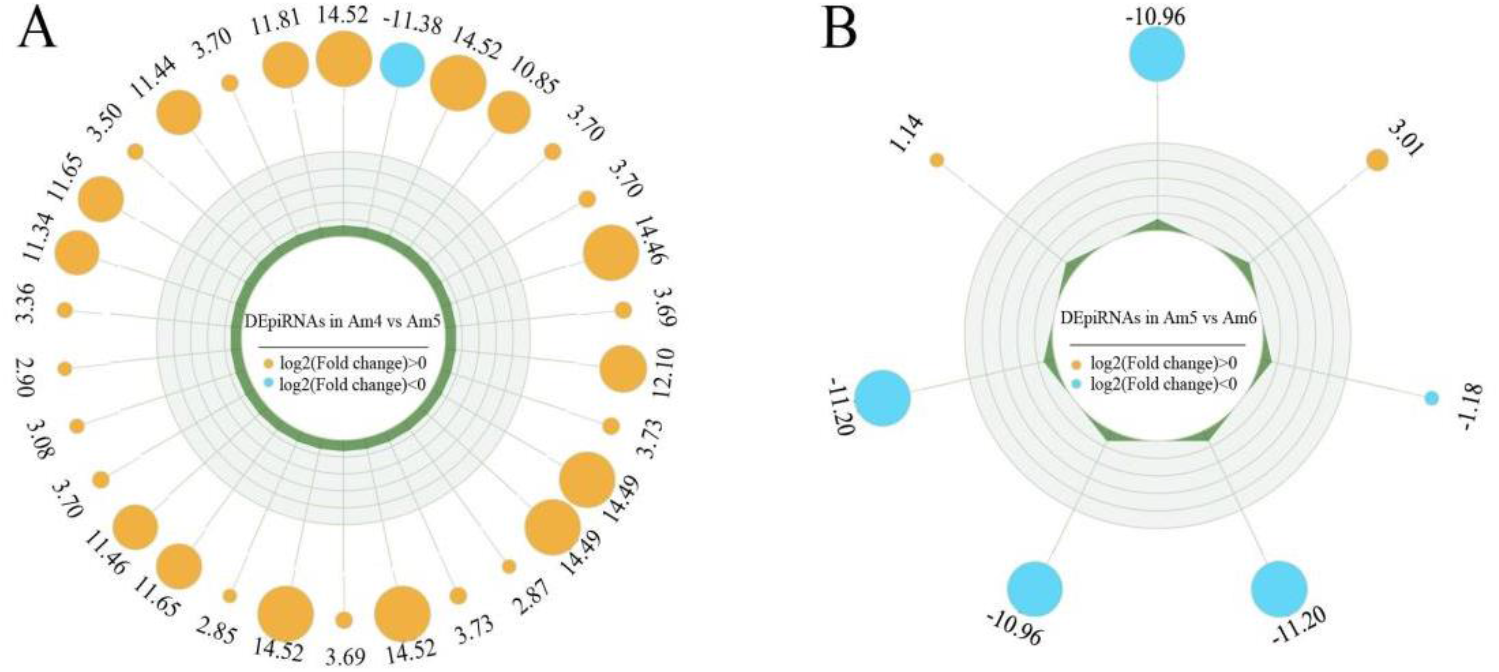
Radar maps of DEpiRNAs. (**A**) DEpiRNAs in Am4 vs. Am5 comparison groups. (**B**) DEpiRNAs in Am5 vs. Am6 comparison groups.

### 3.3. Target prediction and annotation of DEpiRNA

DEpiRNAs in the Am4 vs. Am5 comparison group can target 9, 983 genes, which could be annotated to 20 biological-process-related GO terms such as cellular process and metabolic process, 11 molecular function-related GO terms such as binding and catalytic activity, and 19 cellular component-related GO terms such as cell and membrane part (**Figure 3**A). DEpiRNAs in the Am5 vs. Am6 comparison group can target 1, 936 genes, and these targets could be annotated to a total of 41 GO terms, including cellular process, catalytic activity, and cells. (**Figure 3**B).

**Figure 3.**
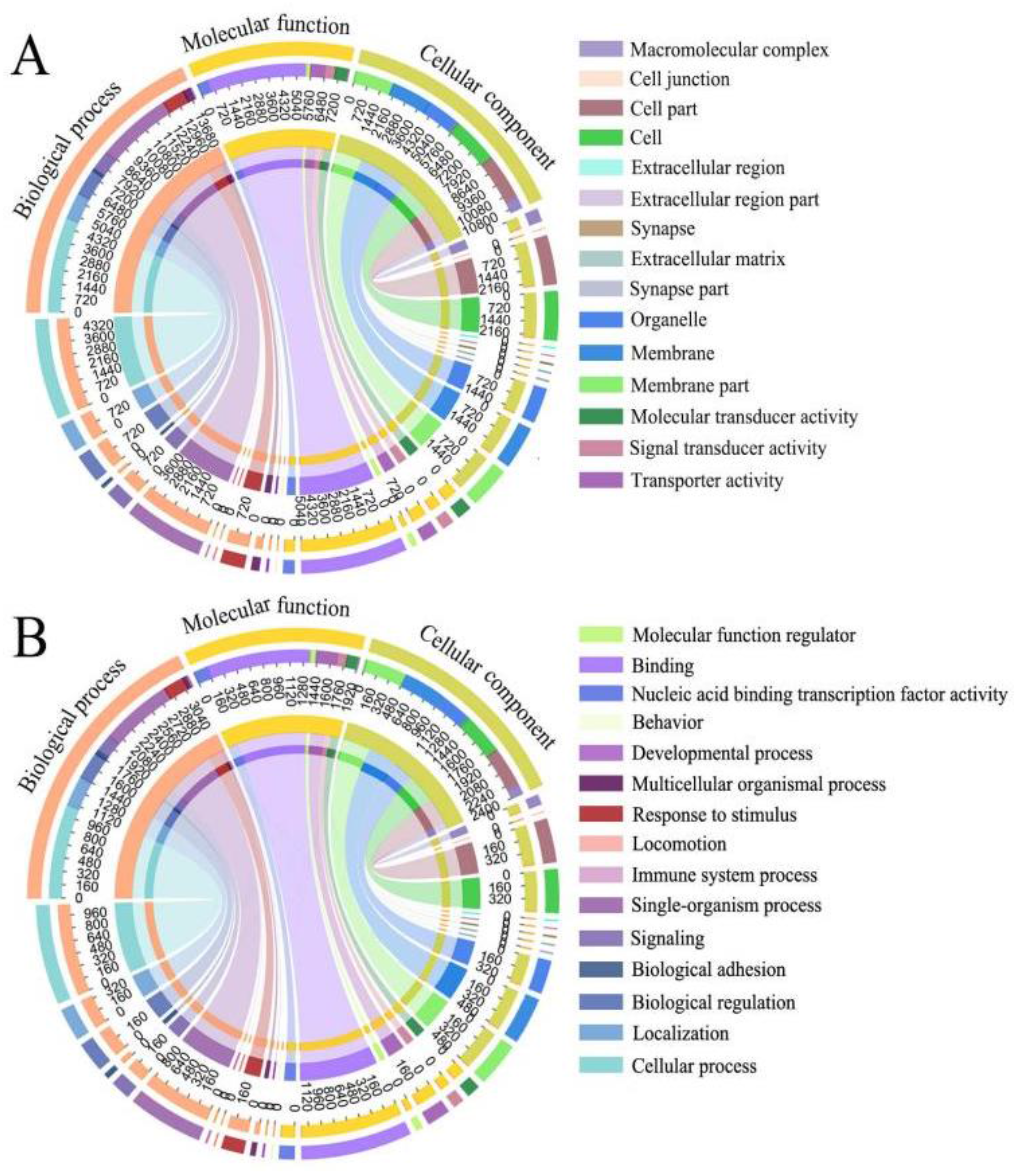
GO database annotation of target genes of DEpiRNAs. (**A**) Targets in Am4 vs. Am5 comparison groups. (**B**) Targets in Am5 vs. Am6 comparison groups.

In addition, target genes of DEpiRNA in Am4 vs. Am5 comparison group could be annotated to 142 pathways such as Wnt signaling pathway, endocytosis, and purine metabolism (**Figure 4**A); those in the Am5 vs. Am6 comparison group can be annotated to 101 pathways including the Hippo signaling pathway, RNA transport, and Lysosome (**Figure 4**B).

**Figure 4.**
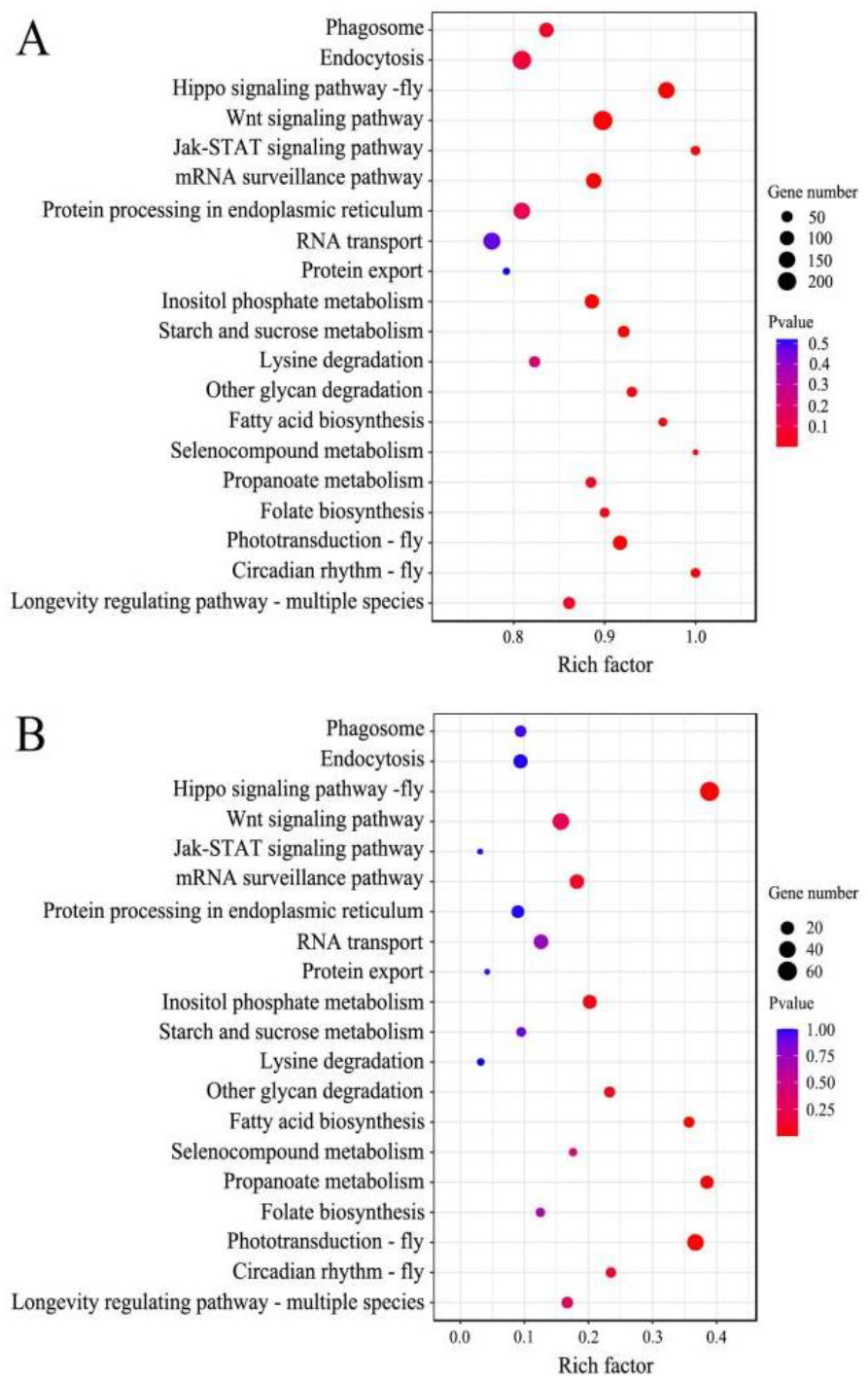
KEGG database annotation of DEpiRNA-targeted genes. (**A**) Targets in Am4 vs. Am5 comparison groups. (**B**) Targets in Am5 vs. Am6 comparison groups.

### 3.4. Investigation of regulatory network between DEpiRNAs and target genes

In the Am4 vs. Am5 comparison group, 54 up-regulated piRNAs could target 9, 398 genes, while 14 down-regulated piRNAs could target 3, 606 genes; each of these DEpiRNAs can target more than two genes, with piR-ame-742536 and piR-ame-856650 binding to the highest number of target genes (1, 421 and 1, 437). Additionally, two up-regulated piRNAs in the Am5 vs. Am6 comparison group could target 604 genes, whereas four down-regulated piRNAs could target 1, 473 genes; each of these DEpiRNAs can target more than two genes, with piR-ame-592661 and piR-ame-31653 linking to the highest number of target genes (447 and 839).

The regulatory network was constructed and analyzed, and the result showed that 202 and 58 target genes in the above-mentioned two comparison groups were involved in seven development-associated signaling pathways such as Hippo, Notch and mTOR signaling pathways, whereas 255 and 39 targets were engaged in seven immune-associated pathways including endocytosis, the Jak/STAT signaling pathway, and ubiquitin-mediated proteolysis (**Figure 5**A). Additionally, 33 and 12 targets were found to be enriched in three energy metabolism pathways, namely sulfur metabolism, nitrogen metabolism, and oxidative phosphorylation (**Figure 5**B).

**Figure 5.**
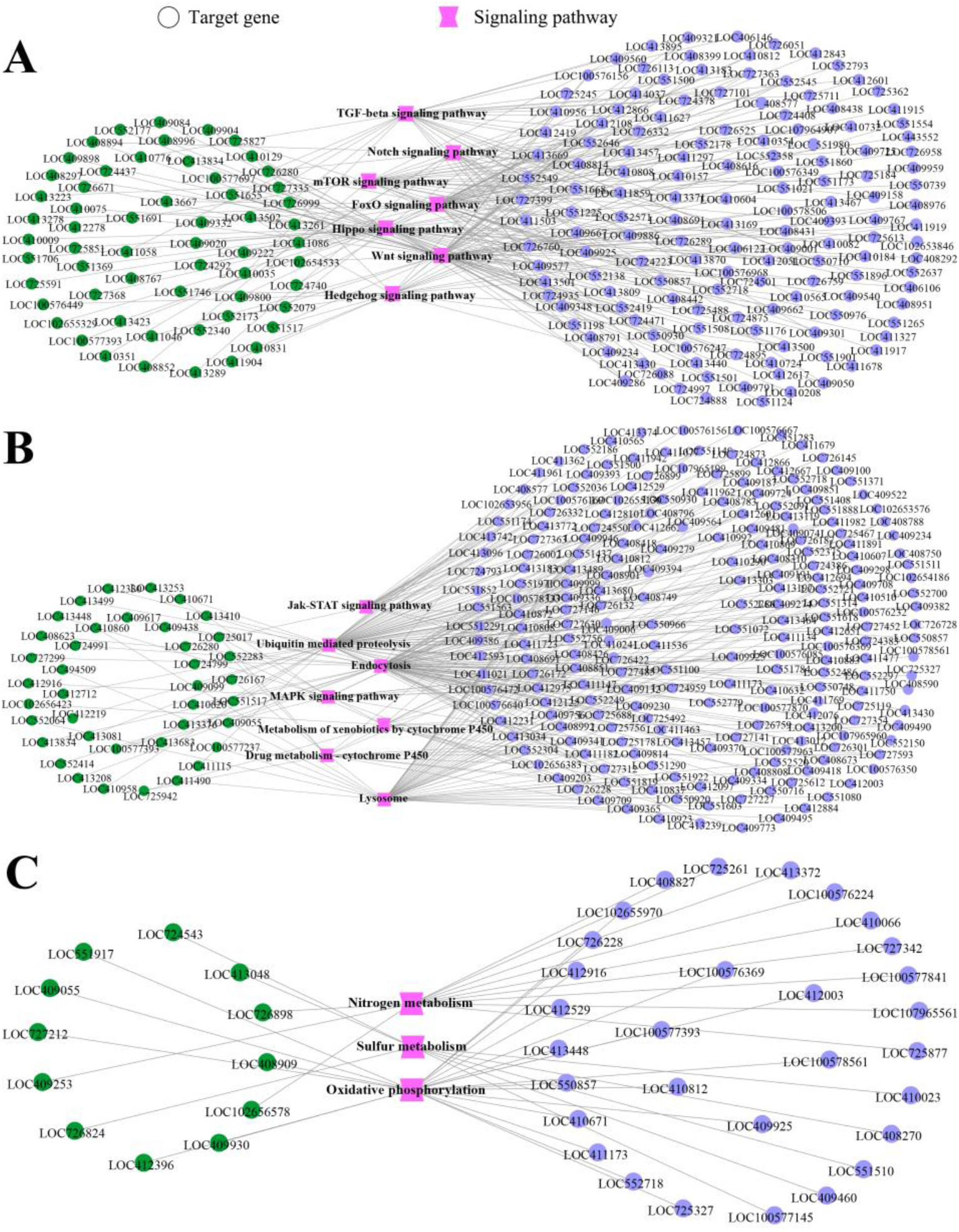
Regulatory network of f DEpiRNAs in Am4 vs. Am5 and Am5 vs. Am6 comparison groups. (**A**) Regulatory network of DEpiRNAs and corresponding targets relative to development-associated pathways. (**B**) Regulatory network of DEpiRNAs and corresponding targets relative to immune-associated pathways. (**C**) Regulatory network of DEpiRNAs and corresponding targets relative to energy metabolism-associated pathways. Purple circles indicate target gene of DEpiRNA in Am4 vs Am5, green circles indicate terget gene of DEpiRNA in Am5 vs Am6, pink concavehexagon indicate signaling paythway.

### 3.5. Stem-loop RT-PCR and RT-qPCR verification of DEpiRNA

The stem-loop RT-PCR result indicated that fragments with expected size (about 80 bp) were amplified from five randomly selected five DEpiRNAs (**Figure 6**), which verified the expression of these DEpiRNAs in the *A. m. ligustica* larval gut.

**Figure 6.**
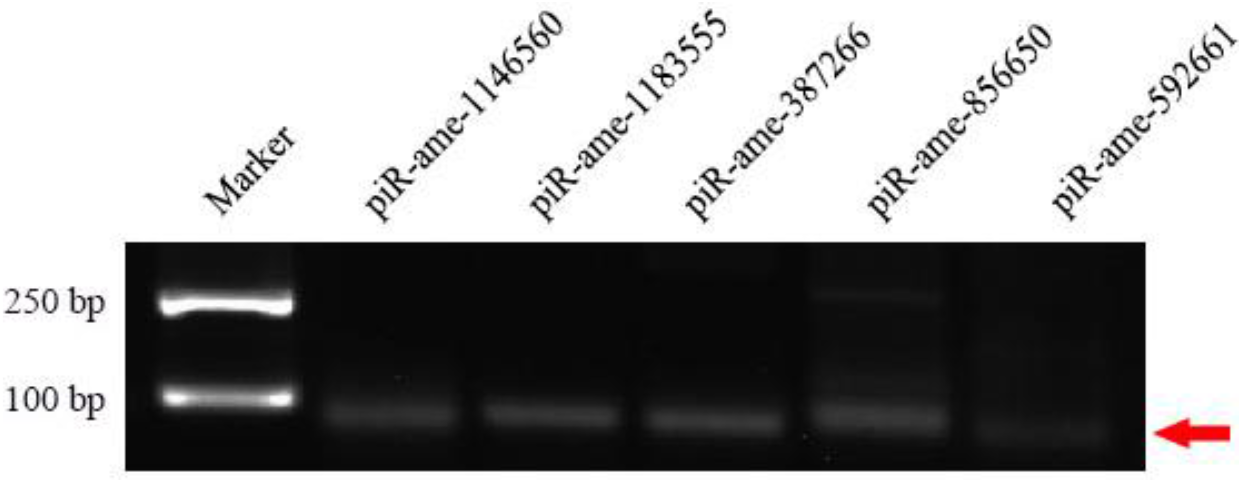
Stem-loop RT-PCR validation of five DEpiRNAs. Red arrow indicates the signal band with expected size.

Further, RT-qPCR result suggested that the expression trend of these five DEpiRNAs were consistent with sRNA-seq datasets, confirming the reliability of our transcriptome data (**Figure 7**).

**Figure 7.**
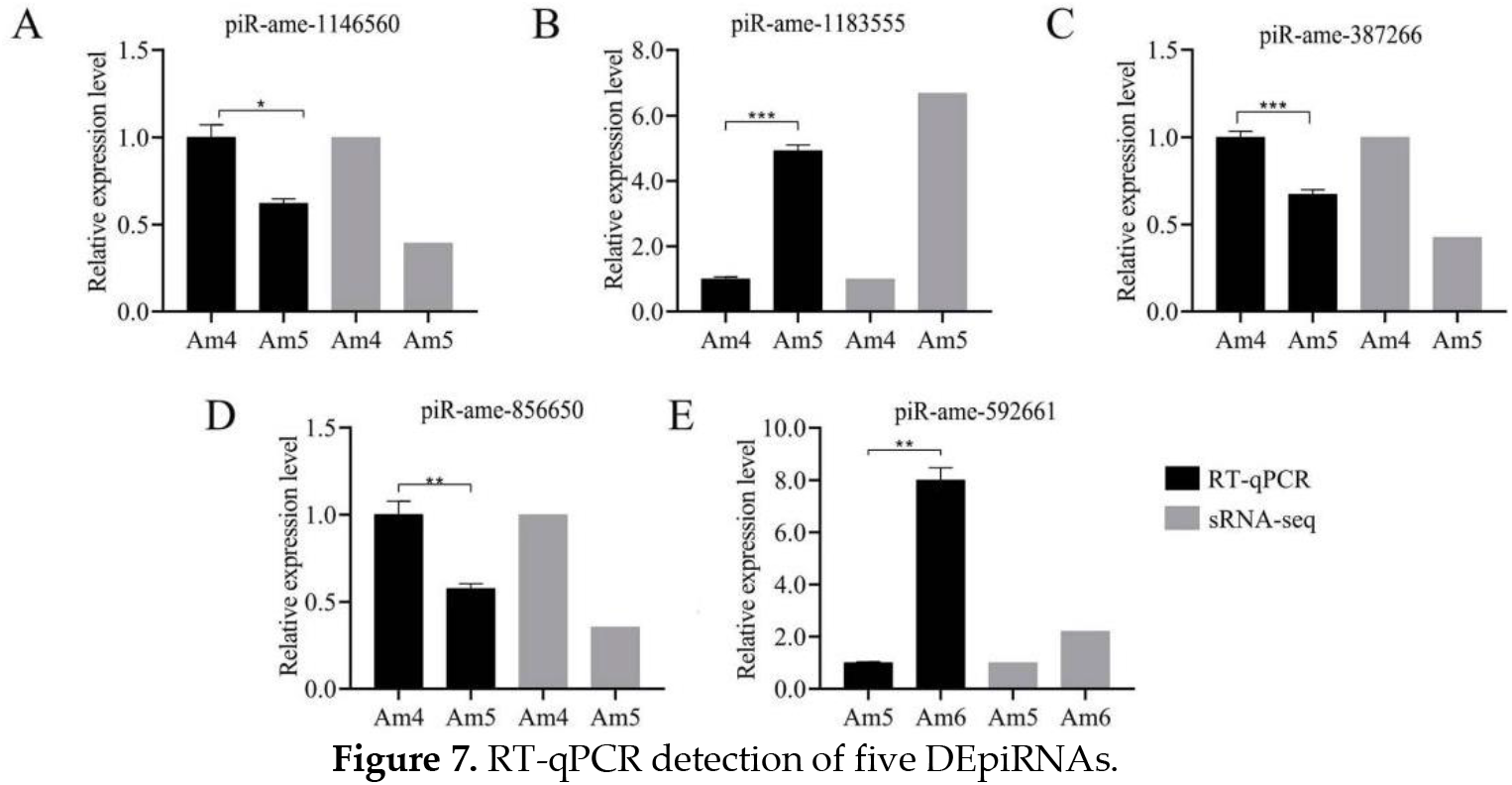
RT-qPCR detection of five DEpiRNAs.

## 4. Discussion

At present, there are few studies on piRNAs in honey bee. Our team previously predicted 596 piRNAs in the *A. m. ligustica* workers’ midguts based on bioinformatics [26]. In this study, 843 piRNAs were identified in the larval guts of *A. m. ligustica*. Further analysis showed that as many as 519 (61.57%) piRNAs were shared by the workers’ midguts and larval guts of *A. m. ligustica*, whereas the numbers of unique piRNAs were 324 and 77. It is inferred that the shared piRNAs are likely to play a fundamental role in various developmental stage of *A. m. ligustica*, while the unique piRNAs may play different roles in different developmental stage. In view of the limited information on bee piRNA, the piRNAs identified in the present study further enrich the reservoir of piRNAs in European honey bee and offer a valuable genetic resource for related studies on other bee species.

In animals, piRNAs were verified to participate in regulation of growth, development, and embryogenesis. Based on overexpression and knockdown of piRNA-3312, Guo et al. [27] found that piRNA-3312 targeted the gut esterase 1 gene to decrease the pyrethroid resistance of *Culex pipiens pallens*. Praher et al. [28] found that piRNAs were significantly differentially expressed in early developmental stages of *Nematostella vectensis*, indicative of the regulatory role of piRNAs in development. Here, 78 and 7 DEpiRNAs were observed in Am4 vs. Am5 and Am5 vs. Am6 comparison groups, respectively, indicating that the process of the larval gut of *A. m. ligustica* was accompanied by differential expression of piRNAs, and these DEpiRNAs may be engaged in regulating development of *A. m. ligustica* larval gut. DEpiRNAs in the Am4 vs. Am5 and Am5 vs. Am6 comparison groups were found to target 9; 983; and 1,936 genes, which were involved in cellular process and localization terms relative to biological processes, membrane parts and membrane terms relative to cellular components, and binding and transporter activity relative to molecular function. Additionally, targets of DEpiRNAs in the aforementioned two comparison groups were involved in four and three development-associated terms such as metabolic process and development process and three and two immune-associated terms such as immune system process and response to stimulus. Targets in these two comparison groups were engaged in 142 and 101 KEGG pathways, including fatty acid metabolism and sulfur metabolism relative to metabolism, RNA polymerase and proteasome relative to genetic information processing, and lysosomes and endocytosis relative to cellular processes. Further analysis indicated that targets were engaged in seven and seven development-related pathways such as the Hippo signaling pathway and Wnt signaling pathway, seven and seven immune-related pathways such as endocytosis and the Jak/STAT signaling pathway, as well as three and three energy metabolism-related pathways such as nitrogen metabolism and sulfur metabolism. These results demonstrate that DEpiRNAs exerted a potential regulatory function in the *A. m. ligustica* larval guts by affecting many biological processes including development, immune defense, and energy metabolism.

Mondal et al. [29] confirmed that piRNA was capable of silencing gene expression in a siRNA-like manner. In the present study, a complex regulatory network between DEpiRNAs and target genes was observed, and all DEpiRNAs had more than two targets, implying that DEpiRNAs may be used by *A. m. ligustica* larvae to modulate target gene expression during gut development. Additionally, piR-ame-742536 (log_2_FC=11.34, *P*=5.42E-07) and piR-ame-856650 (log_2_FC=−1.49, *P*=0.0003) in the Am4 vs. Am5 comparison group could target 1; 421; and 1, 437 genes, respectively, while piR-ame-592661 (log_2_FC=1.14, *P*=0.005) and piR-ame-31653 (log_2_FC=−10.96, *P*=1.34E-05) in the Am5 vs. Am6 comparison group had 447 and 839 target genes, respectively. This indicated that the abovementioned four DEmiRNAs were likely to play crucial parts in the developmental process of *A. m. ligustica* larval guts.

As a highly conserved signaling pathway, the Wnt signaling pathway plays a key role in maintaining the development and homeostasis of animals and in promoting intestinal regeneration [30]. Shah et al. [31] discovered that silencing *Wnt-1* at the larval stage of *Tribolium castaneum* could result the larval death and abnormal pupal and adult development. Fu et al. [32] conducted knockout of the *HaWnt1* gene in *Helicoverpa armigera* using CRISPR/Cas9 technology, and the result showed that the HaWnt1 signaling pathway was essential for embryonic development of *H. armigera*. Here, 68 DEpiRNAs in the Am4 vs. Am5 comparison group could target 58 genes involving the Wnt signaling pathway, including piR-ame-247619 (log_2_FC=2.01, *P*=0.002), piR-ame-750627 (log_2_FC=2.20, *P*=0.003), and piR-ame-990954 (log_2_FC=11.81, *P*=5.82E-05). Additionally, seven DEpiRNAs in the Am5 vs. Am6 comparison group could target 15 Wnt signaling pathway-relevant genes, including piR-ame-1173337 (log_2_FC=−10.96, *P*=1.34E-05), piR-ame-31653 (log_2_FC=−10.96, *P*=1.34E-05), and piR-ame-260979 (log_2_FC=−11.20, *P*=3.78E-05). In insects, the Hippo signaling pathway participates in regulating body size as well as normal growth and development [33]. The *BmSd* gene was characterized as one of the Hippo signaling-pathway-related genes [34]. Yin et al. [34] detected that as many as 57.9% of *Bombyx mori* showed deformed wings after *BmSd* knockdown. Here, we found that 68 DEpiRNAs in Am4 vs. Am5 such as piR-ame-456655 (log_2_FC=14.52, *P*=7.22E-05), piR-ame-5 (log_2_FC=14.52, *P*=7.22E-05), and 14055 (log_2_FC=14.52, *P*=7.22E-05) could target 52 genes involving the Hippo signaling pathway, whereas five DEpiRNAs in Am5 vs. Am6 could target 22 genes involving the Hippo signaling pathway, including piR-ame-31653 (log_2_FC=−10.96, *P*=1.34E-05), piR-ame-1173337 (log_2_FC=−10.96, *P*=1.34E-05), piR-ame-260979 (log_2_FC=−11.20, *P*=3.78E-05), piR-ame-592661 (log_2_FC=1.14, *P*=0.005), and piR-ame-1246710 (log_2_FC=−1.18, *P*=0.045). Together, these results suggested that corresponding DEpiRNAs may affect Wnt and Hippo signaling pathways through regulating target gene expression, further controlling the growth and development of *A. m. ligustica* larval guts. However, additional work is needed for exploration of the underlying mechanism.

The immune system in insects is composed of the humoral immune system dominated by several signaling pathways such as Imd/Toll, JAK/STAT, JNK, and insulin, and cellular immune system represented by phagocytosis, melanization, autophagy, and apoptosis [35]. The JAK/STAT signaling pathway is not only implicated in regulating cell growth, differentiation, apoptosis, and inflammatory immunity but also participates in gut immunity via modulation of intestinal stem cell proliferation and epithelial cell renewal [35,36]. Here, we observed that the JAK/STAT signaling-pathway-related genes were targeted by 54 DEpiRNAs (piR-ame-742536, piR-ame-1183555, and piR-ame-1233036, etc.) in the Am5 vs. Am6 comparison group and 1 DEpiRNA (piR-ame-1243913) in the Am5 vs. Am6 comparison group, suggestive of the involvement of corresponding DEpiRNAs in regulation of immune defense in the larval guts. *Tomato yellow leaf curlvirus* has been proved to enter whitefly *Bemisia tabaci* midgut epithelial cells through receptor-mediated clathrin-dependent endocytosis [37]. Zhang et al. [38] reported that inhibition of endocytosis induced the proliferation of *Drosophila* intestinal stem cells and massive gut hyperplasia, which further affected intestinal development and lifespan. In this study, endocytosis-associated genes were found to be targeted by 60 DEpiRNAs in the Am4 vs. Am5 comparison group including piR-ame-14055 and piR-ame-456655 and five DEpiRNAs in the Am5 vs. Am6 comparison group it was regulated by such as piR-ame-31653 and piR-ame-1173337, indicating that corresponding DEpiRNAs potentially regulated the endocytosis during the development of larval guts. Together, these results demonstrated that DEpiRNAs as potential regulators participated in the development of *A. m. ligustica* larval guts. More efforts are required to elucidate the regulatory function of these DEpiRNAs.

## 5. Conclusion

Taken together, 843 piRNAs were, for the first time, identified in the *A. m. ligustica* larval guts, and the first base of *A. m. ligustica* piRNAs had a C bias. Seventy-eight piRNAs were differentially expressed in 5-day-old larval gut compared with 4-day-old larval gut, while only seven DEpiRNAs were detected in the 6-day-old larval gut compared with 5-day-old larval gut. Additionally, these DEpiRNAs could target 9; 983; and 1, 936 genes, respectively, which were engaged in 50 and 41 functional terms such as the developmental process and immune system process and 142 and 101 KEGG pathways such as the Wnt signaling pathway and endocytosis. Moreover, some DEpiRNAs may modulate expression of corresponding target genes in the *A. m. ligustica* larval guts, further affecting sulfur metabolism, nitrogen metabolism, oxidative phosphorylation, endocytosis, and ubiquitin-mediated proteolysis as well as Hippo, Notch, mTOR, and Jak/STAT signaling pathways.

## Supporting information

Table S1: Primers used in this study

## Supplementary Materials

The following supporting information can be downloaded at: www.mdpi.com/xxx/s1, Table S1: Primers used in this study.

## Author Contributions

Conceptualization, R.G.; methodology, Y.X. and Q.L.; software, Y.X. and Q.L.; validation, X.F., Y.Y. and K.Z., J.Z, H.Z. and Y.Y.; formal analysis, Y.X. and Q.L.; writing—original draft preparation, Y.X; writing—review and editing, Y.X., Q.L. and G.R.; supervision, R.G., D.C., Z.F., Z.L. and T.J; project administration, R.G. and D.C.; funding acquisition, R.G., D.C., and Q.L. All authors have read and agreed to the published version of the manuscript.

## Institutional Review Board Statement

Not applicable.

## Informed Consent Statement

Not applicable.

## Data Availability Statement

In this section, please provide details regarding where data supporting reported results can be found, including links to publicly archived datasets analyzed or generated during the study. Please refer to suggested Data Availability Statements in section “MDPI Research Data Policies” at https://www.mdpi.com/ethics. If the study did not report any data, you might add “Not applicable” here.

## Acknowledgments

R.G. genuinely appreciates the love from his dear wife and daughter.

## References

1. Brennecke, J.; Aravin, A.A.; Stark, A.; Dus, M.; Kellis, M.; Sachidanandam, R.; Hannon, G.J. Discrete small RNA-generating loci as master regulators of transposon activity in *Drosophila*. Cell 2007, 128, 1089–1103.

2. Czech, B.; Hannon, G.J. One loop to rule them all: The ping-pong cycle and piRNA-guided silencing. Trends Biochem. Sci. 2016, 41, 324–337.

3. Fabry, M.H.; Falconio, F.A.; Joud, F.; Lythgoe, E.K.; Czech, B.; Hannon, G.J. Maternally inherited piRNAs direct transient heterochromatin formation at active transposons during early *Drosophila* embryogenesis. Elife 2021, 10, :e68573.

4. Vagin, V.V.; Sigova, A.; Li, C.; Seitz, H.; Gvozdev, V.; Zamore, P.D. A distinct small RNA pathway silences selfish genetic elements in the germline. Science 2006, 313, 320–324.

5. Yamashiro, H.; Siomi, M.C. PIWI-interacting RNA in *Drosophila*: biogenesis, transposon regulation, and beyond. Chem. Rev. 2018, 118, 4404–4421.

6. Sun, Y.H.; Lee, B.; Li, X.Z. The birth of piRNAs: how mammalian piRNAs are produced, originated, and evolved. Mamm. Genome 2022, 33, 293–311.

7. Shen, E.Z.; Chen, H.; Ozturk, A.R.; Tu, S.; Shirayama, M.; Tang, W.; Ding, Y.H.; Dai, S.Y.; Weng, Z.; Mello, C.C. Identification of piRNA binding sites reveals the Argonaute regulatory landscape of the *C. elegans* germline. Cell 2018, 172, 937–951.

8. Pippadpally, S.; Venkatesh, T. Deciphering piRNA biogenesis through cytoplasmic granules, mitochondria and exosomes. Arch. Biochem. Biophys. 2020, 695, 108597.

9. Joosten, J.; Overheul, G.J.; Van Rij, R.P.; Miesen, P. Endogenous piRNA-guided slicing triggers responder and trailer piRNA production from viral RNA in *Aedes aegypti mosquitoes*. Nucleic Acids Res. 2021, 49, 8886–8899.

10. Iwasaki, Y.W.; Siomi, M.C.; Siomi, H. PIWI-interacting RNA: its biogenesis and functions. Annu. Rev. Biochem. 2015, 84, 405–433.

11. Ishizu, H.; Siomi, H.; Siomi, M.C. Biology of PIWI-interacting RNAs: new insights into biogenesis and function inside and outside of germlines. Genes Dev 2012, 26, 2361–2373.

12. Yang, L.; Ge, Y.; Cheng, D.; Nie, Z.; Lv, Z. Detection of piRNAs in whitespotted bamboo shark liver. Gene 2016, 590, 51–56.

13. Kolliopoulou, A.; Santos, D.; Taning, C.; Wynant, N.; Vanden, B.J.; Smagghe, G.; Swevers, L. PIWI pathway against viruses in insects. Wiley Interdiscip. Rev. RNA 2019, 10, e1555.

14. Wang, Y.; Jin, B.; Liu, P.; Li, J.; Chen, X.; Gu, J. piRNA profiling of Dengue virus type 2-infected Asian tiger mosquito and midgut tissues. Viruses 2018, 10, 213.

15. Khalifa, S.; Elshafiey, E.H.; Shetaia, A.A.; El-Wahed, A.; Algethami, A.F.; Musharraf, S.G.; Alajmi, M.F.; Zhao, C.; Masry, S.; Abdel-Daim, M.M.; et al. Overview of bee pollination and its economic value for crop production. Insects 2021, 12, 688.

16. Stein, K.; Coulibaly, D.; Stenchly, K.; Goetze, D.; Porembski, S.; Lindner, A.; Konat, S.; Linsenmair, E.K. Bee pollination increases yield quantity and quality of cash crops in Burkina Faso, West Africa. Sci. Rep. 2017, 7, 17691.

17. Ma, E.; Zhu, Y.; Liu, Z.; Wei, T.; Wang, P.; Cheng, G. Interaction of viruses with the insect intestine. Annu. Rev. Virol. 2021, 8, 115–131.

18. Li, Z.; Hou, M.; Qiu, Y.; Zhao, B.; Nie, H.; Su, S. Changes in antioxidant enzymes activity and metabolomic profiles in the guts of honey bee (*Apis mellifera*) larvae infected with *Ascosphaera apis*. Insects 2020, 11, 419.

19. Dosch, C.; Manigk, A.; Streicher, T.; Tehel, A.; Paxton, R.J.; Tragust, S. The gut microbiota can provide viral tolerance in the honey bee. Microorganisms 2021, 9, 871.

20. Xiong, C.L.; Du, Y.; Chen, D.F.; Zheng, Y.Z.; Fu, Z.M.; Wang, H.P.; Geng, S.H.; Chen, H.Z.; Zhou, D.D.; Wu, S.Z.; Shi, C.Y.; Guo, R. Bioinformatic prediction and analysis of miRNAs in the *Apis mellifera ligustica* larvae gut. Chin. J. Appl. Entomol. 201855, 1023–1033.

21. Allen, E.; Xie, Z.; Gustafson, A.M.; Carrington, J.C. microRNA-directed phasing during trans-acting siRNA biogenesis in plants. Cell 2005, 121, 207–221.

22. Smoot, M.E.; Ono, K.; Ruscheinski, J.; Wang, P.L.; Ideker, T. Cytoscape 2.8: new features for data integration and network visualization. Bioinformatics 2011, 27, 431–432.

23. Livak, K.J.; Schmittgen, T.D. Analysis of relative gene expression data using real-time quantitative PCR and the 2(-Delta Delta C(T)) Method. Methods 2001, 25, 402–408.

24. Fisher R.A. The use of multiple measurements in taxonomic problems, Ann. Eugen. 1936, 7, 179–188

25. R Core Team. R: A Language and Environment for Statistical Computing; R Foundation for Statistical Computing. Available online: https://www.R-project.org/ (accessed on 3 February 2021).

26. Fan X.X.; Long Q; Sun M.H.; Guo Y.L.; Zhao H.D.; Song Y.M.; Kang Y.X.; Gu X.Y.; Chen D.F.; Guo R. Identification and analysis of piRNAs in *Apis mellifera ligustica* workers’ midguts. Acta. Entomol. Sin. 2022.

27. Guo, J.; Ye, W.; Liu, X.; Sun, X.; Guo, Q.; Huang, Y.; Ma, L.; Sun, Y.; Shen, B.; Zhou, D.; et al. piRNA-3312: A putative role for pyrethroid resistance in *Culex pipiens pallens* (Diptera: Culicidae). J. Med. Entomol. 2017, 54, 1013–1018.

28. Praher, D.; Zimmermann, B.; Genikhovich, G.; Columbus-Shenkar, Y.; Modepalli, V.; Aharoni, R.; Moran, Y.; Technau, U. Characterization of the piRNA pathway during development of the sea anemone *Nematostella vectensis*. RNA Biol. 2017, 14, 1727–1741.

29. Mondal, M.; Brown, J.K.; Flynt, A. Exploiting somatic piRNAs in *Bemisia tabaci* enables novel gene silencing through RNA feeding. Life Sci. Alliance 2020, 3, e202000731.

30. Yuan, J.; Gao, Y.; Sun, L.; Jin, S.; Zhang, X.; Liu, C.; Li, F.; Xiang, J. Wnt signaling pathway linked to intestinal regeneration via evolutionary patterns and gene expression in the sea cucumber *Apostichopus japonicus*. Front. Genet. 2019, 10, 112.

31. Shah, M.V.; Namigai, E.K.; Suzuki, Y. The role of canonical Wnt signaling in leg regeneration and metamorphosis in the red flour beetle *Tribolium castaneum*. Mech. Dev. 2011, 128, 342–358.

32. Fu, X.Z.; Li, R.; Qiu, Q.Q.; Wang, M.K.; Zha, T.; Zhou, L. Study on the function of *Helicoverpa armigera Wnt1* gene using CRISPR/Cas9 system. J. Asia Pac. Entomol. 2022, 25, 101869.

33. Xu, X.; Bi, H.L.; Zhang, Z.J.; Yang, Y.; Li, K.; Huang, Y.P.; Zhang, Y.; He, L. BmHpo mutation induces smaller body size and late stage larvae lethality in the silkworm, *Bombyx mori*. Insect Sci. 2018, 25, 1006–1016.

34. Yin, J.; Zhang, J.; Li, T.; Sun, X.; Qin, S.; Hou, C.X.; Zhang, G.Z.; Li, M.W. BmSd gene regulates the silkworm wing size by affecting the Hippo pathway. Insect Sci. 2020, 27, 655–664.

35. Bang, I.S. JAK/STAT signaling in insect innate immunity. Entomol. Res. 2019, 49, 339–353.

36. Xin, P.; Xu, X.; Deng, C.; Liu, S.; Wang, Y.; Zhou, X.; Ma, H.; Wei, D.; Sun, S. The role of JAK/STAT signaling pathway and its inhibitors in diseases. Int. Immunopharmacol. 2020, 80, 106210.

37. Pan, L.L.; Chen, Q.F.; Zhao, J.J.; Guo, T.; Wang, X.W.; Hariton-Shalev, A.; Czosnek, H.; Liu, S.S. Clathrin-mediated endocytosis is involved in Tomato yellow leaf curl virus transport across the midgut barrier of its whitefly vector. Virology 2017, 502, 152–159.

38. Zhang, P.; Holowatyj, A.N.; Ulrich, C.M.; Edgar, B.A. Tumor suppressive autophagy in intestinal stem cells controls gut homeostasis. Autophagy 2019, 15, 1668–1670.

